# Optimization of MR-ARFI for Human Transcranial Focused Ultrasound

**DOI:** 10.1101/2024.11.13.623314

**Authors:** Morteza Mohammadjavadi, Ryan T. Ash, Gary H. Glover, Kim Butts Pauly

## Abstract

Magnetic resonance acoustic radiation force imaging (MR-ARFI) is an exceptionally promising technique to non-invasively confirm targeting accuracy and estimate exposure of low-intensity transcranial focused ultrasound stimulation. MR-ARFI uses magnetic field motion encoding gradients to visualize the MR phase changes generated by microscopic displacements at the ultrasound focus. Implementing MR-ARFI in the human central nervous system has been hindered by 1) phase distortion caused by subject motion, and 2) insufficient signal-to-noise ratio at low (<1.0 MPa) ultrasound pressures. The purpose of this study was to optimize human MR-ARFI to allow reduced ultrasound exposure while at the same time being robust to bulk and physiological motion. We demonstrate that a time series of single-shot spiral acquisitions, while triggering ultrasound on and off in blocks, provides ARFI maps that with correction are largely immune to bulk and pulsatile brain motion. Furthermore, the time series approach allows for a reduction in ultrasound exposure per slice while improving motion robustness with reduced scan time. The focused ultrasound beam can be visualized in an 80 second scan with our protocol, enabling iteration for image-guided targeting. We demonstrated robust ARFI signals at the expected target in 4 participants. Our results provide persuasive proof-of-principle that MR-ARFI can be used as a tool to guide ultrasound-based precision neural circuit therapeutics.

## Introduction

MR-ARFI is a powerful method to confirm precise anatomical targeting of focused ultrasound for therapeutic applications^1^. As opposed to high-intensity focused ultrasound ablation procedures, in which an increase in temperature can be visualized with MR-thermometry^2–4^, in low-intensity applications such as neuromodulation, targeted drug or other payload release, and blood-brain barrier opening, the temperature rise is generally undetectable by MR^5–7^. The need for a technique such as ARFI has become more important since the emergence of transcranial ultrasound applications, which implements transcranial focused ultrasound as a tool for noninvasive brain stimulation, blood brain barrier (BBB) opening or ablation. Noninvasive transcranial ultrasound holds great promise for non-invasive treatment of brain disorders and causal manipulation in human neuroscience, but the accuracy and focality of transcranial ultrasound are severely impacted by refraction, reflection, and attenuation of ultrasound waves by the skull. Because the ultrasound focus is small (<1 cm), and the targets are often deep inside the brain (usually >6 cm), even small deviations (5-10°) in the incident angle of the beam as it passes through the skull can lead to completely missing the target. Simulations greatly help to correct for skull aberrations but are limited by uncertainty in assigning acoustic properties to skull images, simplifications inherent in computational modeling, and various experimental factors not included in the simulations. Therefore, without a ground-truth measure of the focus, the reliability and consistency of transcranial ultrasound will remain unknown.

MR-ARFI measures the phase perturbations induced by ultrasound waves in a motion-encoding magnetic field gradient. MR-ARFI has been shown to reliably visualize the ultrasound focus in large animal models *in vivo*, but the 2DFT acquisitions that were previously employed required relatively high acoustic pressures (2-3 MPa)^8–11^ in order to overcome physiological motion artifacts. In this prior work, two 2DFT spin echo pulse acquisitions were utilized to acquire the complex images, from which the phase difference is extracted to create tissue displacement (ARFI) maps^12^. These could be either ultrasound-on/ultrasound-off with the same encoding gradients or ultrasound-on with encoding gradients inverted in the second acquisition^13^. Either way, while the reconstructed magnitude images were essentially immune to small bulk motions in 2DFT, the corresponding phase images were degraded by bulk motion and pulsatile brain artifacts^14,15^. This is true because ultrasound-induced tissue displacement is only a few microns, compared to bulk motion that can be substantially larger in human scans.

An alternative is to use single-shot acquisitions, with either EPI or spiral imaging, to generate the two complex images^16^, which may have reduced motion artifacts. However, here we use a timeseries of singe-shot spiral images, while triggering the ultrasound on and off in blocks, analogous to fMRI block task designs^17^ and compare ARFI results with those of the 2DFT design. Induced pseudo-random motion of a phantom allows accurate comparison of the motion sensitivity of the two methods.

## Methods and Materials

### Magnetic resonance imaging

MR images were acquired on a 3T scanner (DV750, GE Healthcare, Milwaukee, WI) with spin echo (SE) sequences. Inverted bipolar motion encoding gradients (MEGs) were employed, with a total duration of 20 or 30ms, corresponding diffusion b-values^15^ of 29.5 or 100.5 s/mm^2^ and amplitude of 50 mT/m. An 8-channel head coil (MRI Devices) was used.

### 2DFT ARFI Image acquisition

For comparison with the proposed timeseries method, we used a conventional 2DFT SE sequence with TR/TE of 800ms/40ms, resolution of 0.86 × 1.72 mm, 4 mm slices, scan time of 80 seconds. Two separate scans were obtained, without and with ultrasound, for subsequent processing of the complex images to obtain the phase difference caused by ARF. The longer scan time (240s total for two scans vs. 80s for the single spiral scan) approximately compensated SNR for the increased spatial resolution of the 2DFT sequence.

### MR-ARFI using Spiral Time series method

Introduced in this study as an alternative to conventional 2DFT imaging, we utilized a timeseries approach with a variable-density single-shot spiral-out MRI sequence, acquiring 100 time frames with TR/TE = 800ms/40-50ms, 3 slices of 4 or 5 mm thick, 2.75 × 2.75 transverse resolution, reconstructed into 1.72 × 1.72 mm pixel size (128 × 128 matrix), scan time 80 seconds. A block design analogous to that used in fMRI^17,18^ is used to turn on and off the ultrasound during the scan in blocks of 25 time frames, as shown in Fig. 1A. The variable-density spiral readout duration is 16.9 ms, less than half that of a comparable resolution single-shot EPI trajectory (40.2ms), which thereby provides greater immunity to motion^19^.The alternating on-off blocks allowed detrending of the magnitude and phase images, to correct for drift.

**Figure 1.**
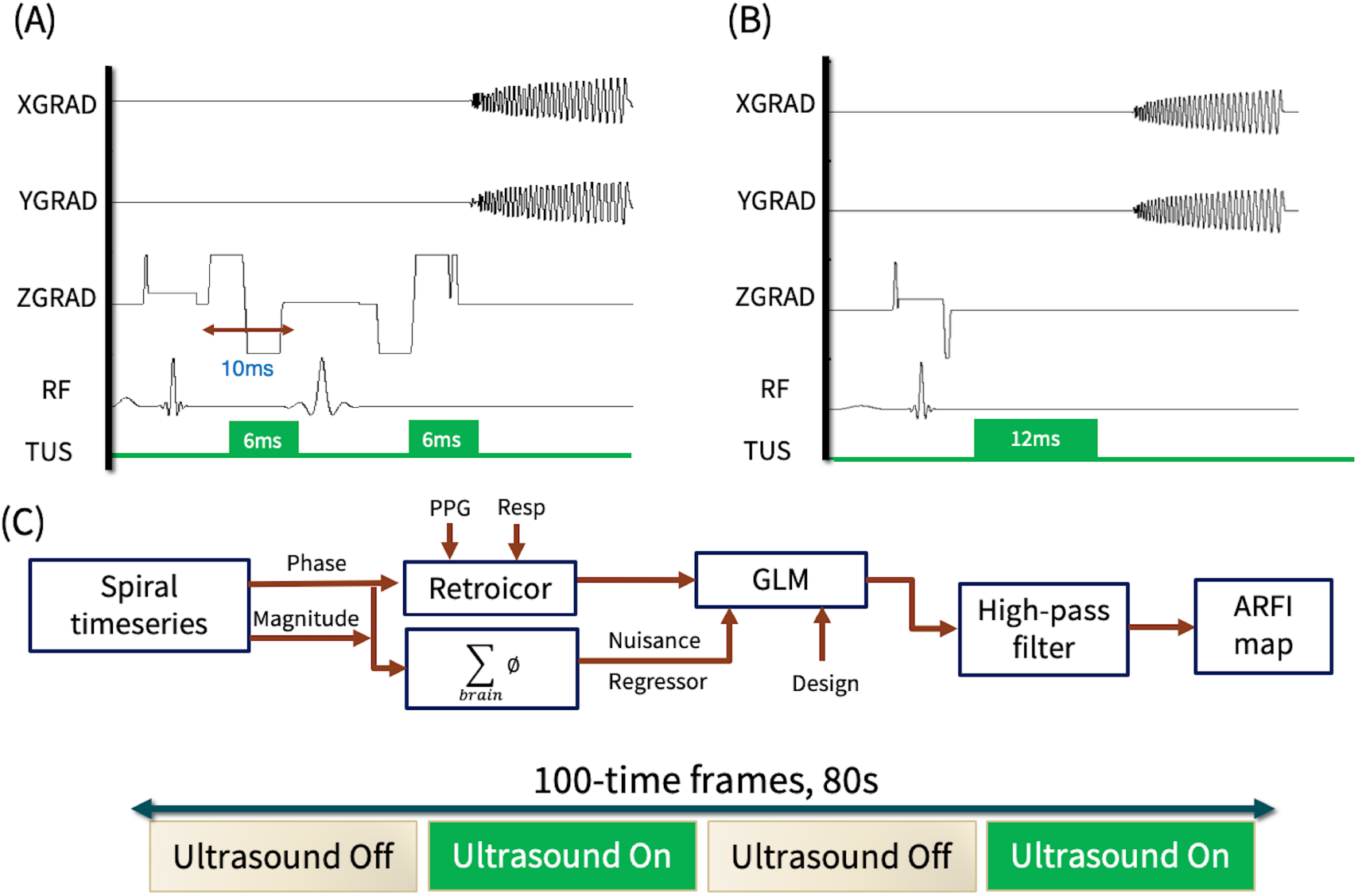
Pulse sequence and processing pipeline. (A) single-shot (SS) spiral ARFI spin echo pulse sequence, showing repeated bipolar MEGs together with ultrasound pulses to generate phase encoding of tissue displacement. The sequence is repeated an equal number of times with ultrasound on and with ultrasound off. (B) The thermometry gradient echo sequence using the same duration of ultrasound as used for ARFI. (C) The processing pipeline for producing ARFI statistical map using SS spiral pulse sequence, and the 80-second block-design protocol for measuring ARF.

During MR-ARFI, focused ultrasound was applied during the second lobe of the first bipolar and the second lobe of the second inverted bipolar motion encoding gradient (Fig. 1A). The resulting phase shift Δ*ϕ* of tissue displaced by the sonication is

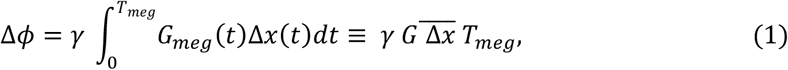

where γ is the gyromagnetic ratio for protons, G_meg_(t) is the MEG waveform, Δx(t) is the time-dependent displacement, 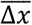 is defined as an average displacement, G is the maximum amplitude of the MEG, and T_meg_ is the total duration of the MEGs.

### Generation of ARFI maps

The spiral acquisitions were reconstructed as a complex (magnitude & phase) timeseries. During reconstruction, the phase of the second time frame was subtracted from all subsequent time frames to keep the phase timeseries well within ±π, thereby avoiding phase wraparound. With reference to Fig. 1A, the pixel-wise average of the magnitude timeseries was used to generate a mask of those pixels with magnitude greater than 25% of the peak brain signal. This mask was applied to the phase timeseries to exclude non-brain pixels during subsequent processing with a General Linear Model (GLM) to identify voxels with significant phase differences between ultrasound-on/off blocks.

### Bulk motion correction

To correct for head motion in the longitudinal direction (direction of sonication beam), the average phase of the brain slices collected was calculated for every time frame, as a regressor of no interest in the general linear model. Since the sonication region was a tiny fraction of the brain volume, this average signal demonstrated bulk motion of the brain, but was minimally affected by ARF(t) modulation.

### Physiological correction

In fMRI, it is standard practice to correct timeseries data for quasiperiodic physiological artifacts^20^. These include cardiovascular-induced brain pulsatility (monitored by photoplethysmography [PPG] on a finger), respiratory-induced changes in blood volume, and resonance frequency shifts caused by diaphragm movement (monitored with a respiration belt). For the purpose of removing physiological noise, we applied RETROICOR^21^ to the phase timeseries which corrects for cardiac and respiratory-induced phase artifacts (Fig. 1A).

### General Linear Model (GLM)

The corrected timeseries of phase images Δ∅(*t*) was statistically tested for the inclusion of signals that matched the off-on-off-on block design *d(t)* of the ultrasound stimulation (Fig. 1C) using a GLM^18,22^. The GLM is defined as

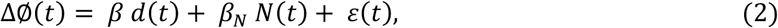

where *β* is the amplitude of the design contained in the measured timeseries data for every voxel Δ∅(*t*), N(t) is a nuisance regressor of no interest with *β*_*N*_ its amplitude, and *ε*(*t*) is residual noise. Then *β* represents the observed phase change due to ARF.

The average phase timeseries is included as a nuisance regressor to diminish bulk motion effects, and a quadratic detrending regressor is included to diminish slow phase drifts in the timeseries. The resulting Statistical Parametric Maps (SPMs)were presented as T-scores that depict probability that the block design was observed in the timeseries.

As the last step in the processing pipeline (Fig 1C), spatial filtering was employed to reduce residual low-spatial frequency bias in the SPMs. After 2DFT of the maps, the Fourier components within a radius of 2 of the k-space center were nulled, and the inverse transform applied. This high-pass filter minimally affected the amplitude of the ARFI-voxels (98% retention) because of the spatially discrete nature of ARFI.

### Phantom ARFI

To optimize the design for tissue response with MR-ARFI and examine sensitivity to bulk motion, a 1.0 MHz 128-element phased array US transducer (Image Guided Therapy, Paris, France) was focused to a depth of 60 mm to generate micro-displacement in a soft tissue-mimicking phantom (Computerized Imaging Referencing System, Virginia, USA). To generate controlled motion, the phantom with transducer attached was placed in the head coil on a platform with rollers that was attached by a 2.3m rod to a stepper-motor translation stage at the end of the patient table. The motor was driven in pseudo-random steps of multiples of 25.4 micrometers (microns) asynchronously with the ARFI scan TR.

As part of the design optimization, it was useful to estimate the ARF time constant. In Kaye et al, 2013, rise time response of pig brain post-mortem was measured to be 5.9ms. The same time-constant value was used as a reference to simulate encoded phase with inverted bipolar gradient. Time delays of 1ms and 2ms were applied between the ultrasound pulse and the MEGs for a range of time constant to evaluate the simulated encoded phase. To validate the simulation values, we performed ARFI experiments using our phantom with similar properties to soft tissue (Young’s Modulus: 12 +/-2 kPa, Speed of sound: 1540 +/-10 m/s, Attenuation coefficient: 0.5 +/-0.1 dB/cm/MHz).

### Thermometry Pulse sequence

To investigate if there was any temperature increase from the sonication, MR-thermometry was obtained using the proton resonant frequency shift method, with a thermal coefficient of 0.01 ppm/°C^23^. A gradient-recalled echo (GRE) pulse sequence otherwise identical to the single-shot spiral SE sequence employed for ARFI was used to collect a timeseries of 100 time frames, while the ultrasound was applied with identical duration to that with ARFI in a block design. The motion encoding gradients were omitted for improved SNR (eliminating diffusion losses and shortening TE) because they did not affect the heat deposition (Fig 1B). A TR/TE of 800/40 ms was used, with identical slice positioning as for the ARFI acquisition. The same processing of the complex timeseries data was used to generate maps of temperature rise.

### Transcranial Ultrasound in Human

A 500 kHz 4 element annular array (NeuroFUS System, UK) was used with a free-water peak pressure of approximately 1.48 MPa, as measured with a calibrated hydrophone (Onda, Sunnyvale, USA). During spiral scans, two 6 ms pulses with a free field value of 1.48 MPa (*I*_*sppa*_ = 68 *W*⁄*cm*^2^) at 60 mm depth were applied and repeated in on-off blocks of 25 time frames (Fig. 1A). Using conservative derating as described by Attali et al.^24^ for the skull and 0.5dB/cm/MHz for brain tissue, this yielded an estimate of pressure of 0.83 MPa and a MI_tc_ of 1.17^25^ and a total exposure of 38 *J*⁄*cm*^2^. Individual simulations from the four subjects gave peak pressure estimates from 500-800 kPa depending on the participants’ skull properties (see simulations below), with MI_tc_ values of 0.7 – 1.13. Ultrasound gel (Aquasonic 100 Ultrasonic gel, Parker Laboratories, NJ, USA) and a 10 mm thick gel-pad (Aquaflex, Parker Laboratories, NJ, USA) were used to couple the transducer to the participants’ temporal bone. No specific target was identified for sonication. However, a 60 mm depth of focus was used, and the transducer was applied to the temporal bone, consistently across the subjects. The details of the ultrasound parameters are provided in the supplemental materials. This study was approved by the Institutional Review Board of Stanford University. All subjects provided informed consent.

### Acoustic Pressure and Temperature Simulation

Simulations were used to estimate the ultrasound pressure and the temperature rise at the focus. Biophysical head models were generated with the SimNIBS Charm function^26^ using each participant’s T1 and T2 scans. The transducer position was determined in Brainsight software (Rogue Research Inc., CAN) using the planning scans from each experiment. The ultrasound beam was then simulated in open-source BabelBrain software^27^ using each participant’s SIMNIBS head model and ZTE pseudo-CT skull image^28^. BabelBrain generates 3D volumes of the estimated intensity and temperature rise at each pixel in the beam path.

## Results

### ARF time constant simulation and measurements

We first used simulations to parameterize how the brain ARF time constant and ultrasound-MEG time delays influence ARFI measurements (quantified as the encoded phase). The ARF time constant for cadaverous pig brain was quantified as ∼6 ms in Kaye et., al 2013^14^. Figure 2A shows example simulation results, depicting the simulated encoded phase generated by two 6-ms ultrasound pulses at the measured 6 ms time constant, with a 0 ms, 1 ms and 2 ms time-delay between ultrasound pulse and the MEG. This figure shows greater simulated phase using 2 ms time-delay. We then simulated ARFI measurements for a range of ARF time constants and ultrasound-MEG delays. Figure 2B shows simulation results estimating encoded phase for a range of time constant values and two different time delays applied between ultrasound and MEGs. As can be seen, for time constant values greater than 5 ms, a 2 ms time shift gives the greater encoded phase. We then tested the impact of the ultrasound-MEG time delay on measured encoded phase in a CIRS phantom (Fig. 2C,D). Figure 2C shows MR-ARFI data in CIRS phantom at 1 ms (left) and 2 ms (right) ultrasound-MEG time delays. The ARFI value (t-score) was higher at a 2 ms delay compared to a 1 ms delay (Fig. 3D), confirming our simulation results. Therefore, a ultrasound-MEG time delay of 2 ms was employed to maximize displacement in subsequent reported experiments.

**Figure 2.**
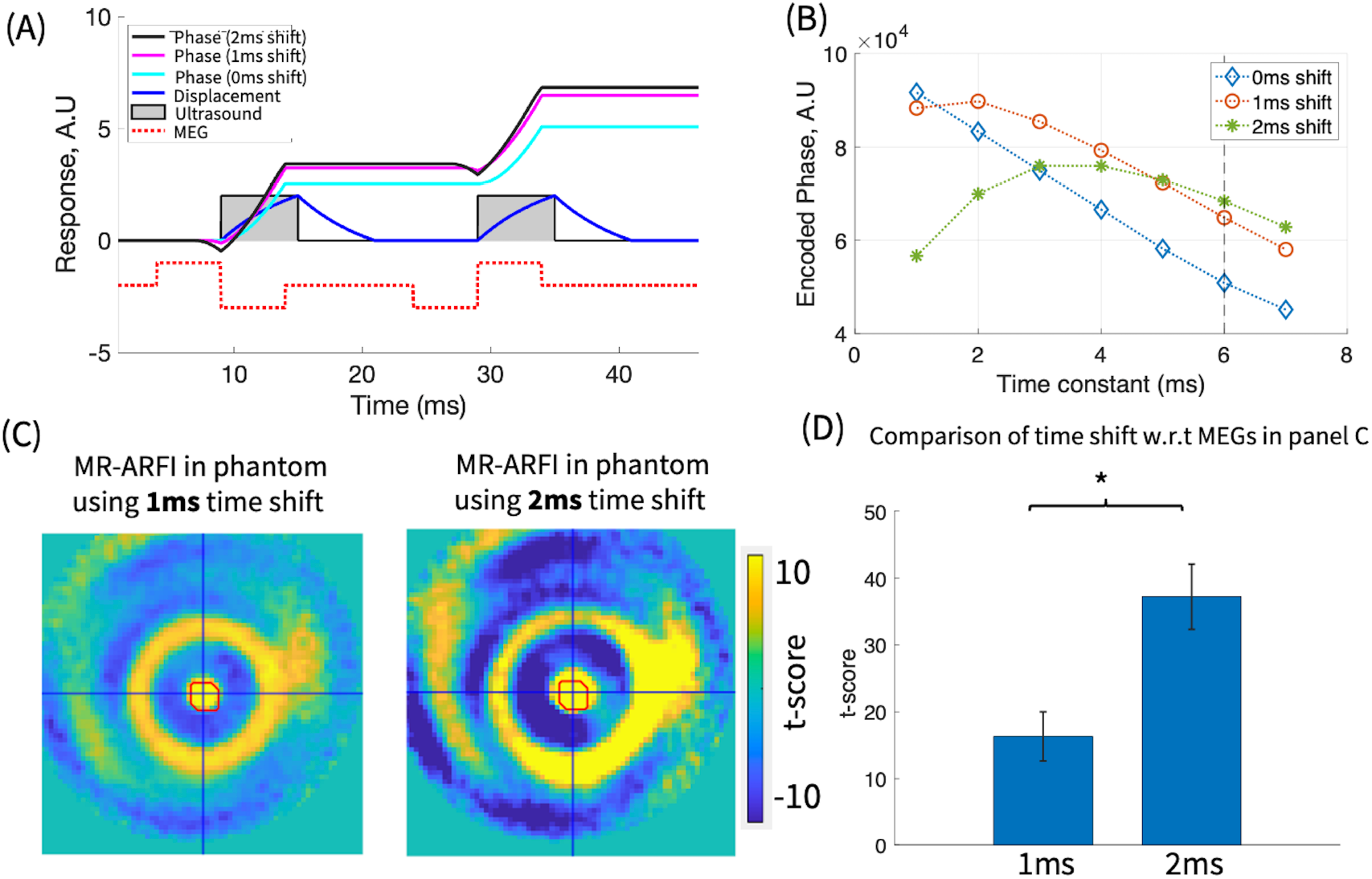
Phantom acoustic radiation force (ARF) time constant measurement and benefit of ultrasound delay. (A) Simulation of encoded phase for ARF time constant of 6 ms using inverted bipolar gradients. The simulated phase is shown in cyan, pink and black color for delaying ultrasound w.r.t MEGs at 0ms, 1ms and 2ms, respectively. (B) Simulated encoded phase for a range of time constant values from1 to 7ms. This figure shows that applying 2ms delays to ultrasound relative to MEGs resulted in a higher phase when the time constant is more than 5ms. (C) Experimental MR-ARFI data in a tissue-mimicking phantom in the middle slice using 1ms (left) and 2ms (right) time shift. The red circle area is defined by a matrix of 3×3 around the maximum value to calculate average t-score. (D) MR-ARFI signal (t-score value) was doubled with 2-ms ultrasound-MEG delay compared to a 1-ms delay (error bars show SEM over three slices).

**Figure 3.**
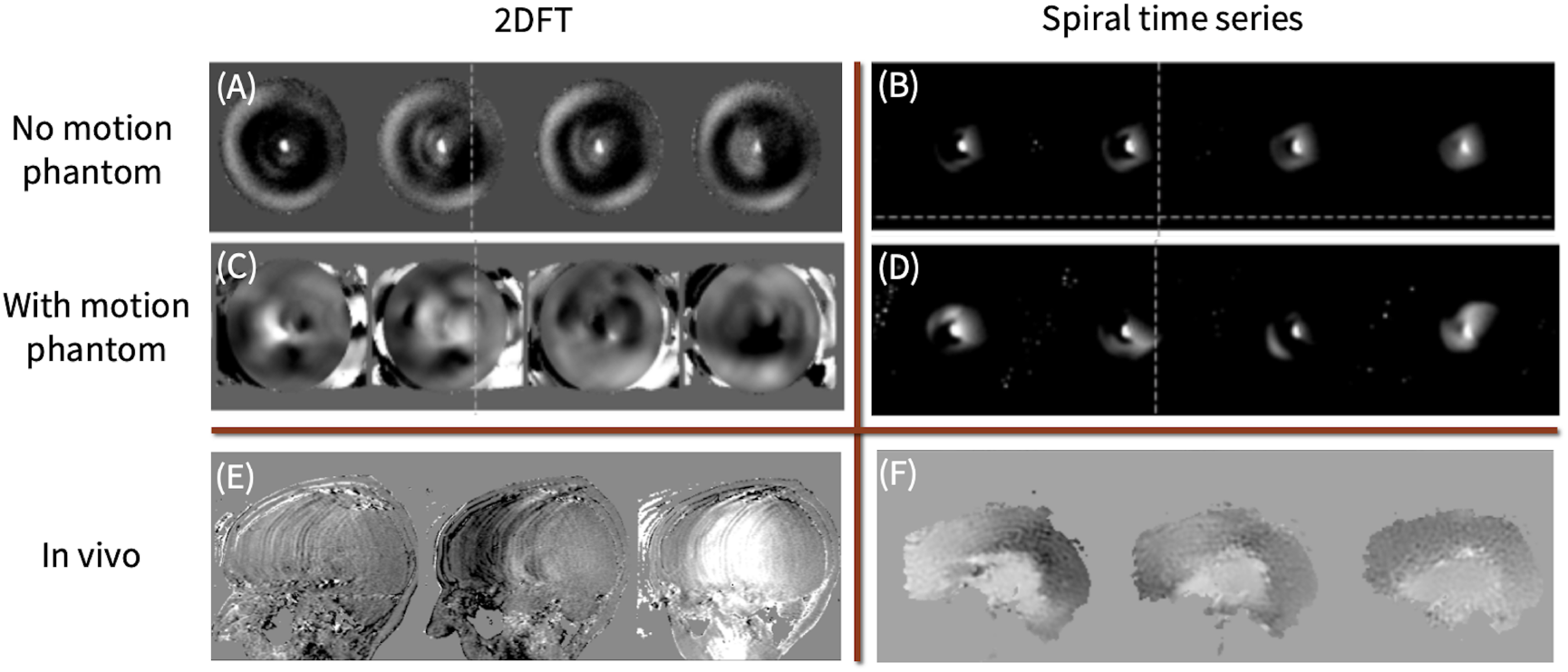
Motion correction. ARFI images obtained with (A) 2DFT, no motion in phantom; (B) Single-shot timeseries spiral, no motion in phantom; (C) 2DFT, the focal spot with ±75 microns random motion is heavily distorted; (D) Single-shot timeseries spiral, with the same motion, but the spiral method retains utility. Phase images obtained using 2DFT MR-ARFI pulse sequence in (E) and parametric maps of single shot spiral timeseries shown in (F).

### Motion Correction using Time series spiral method

We then quantified how motion influences ARFI signal in phantoms. Figure 3 shows ARFI maps acquired with (A) 2DFT, no motion; (B) spiral timeseries, no motion; (C) 2DFT with ±75 micron random motion; (D) spiral timeseries with the same motion. As may be seen, the two methods are roughly equivalent with no intentional motion, but the 2DFT ARFI maps with even the small motion applied fail to show a detectable focal spot, while with the spiral maps the focal spot remains detectable. Panels E shows 2DFT in human in which motion artifact distorted the phase image whereas in panel F we demonstrated motion-robust spiral timeseries. It is notable that the spiral scan time was only 40% that of the 2DFT acquisition, suggesting that even better ARFI could be obtained in the presence of motion, if the same scan time had been used.

### Acoustic and Bioheat simulation comparison to MR-thermometry in the brain

As a last step in our analysis, we wanted to determine if our protocol would generate any significant tissue heating. We performed realistic bioheat simulations with Babelbrain for each participant using our ARFI protocol as input. Fig. 4A,B depicts the simulation results for the subject with the least attenuative skull, showing a maximum in-brain intensity of 21 W/cm^2^ (panel A) and a maximum in-brain temperature rise of 0.5°C (panel B). Next, we looked at the MR-thermometry measures of tissue heating, which were acquired during sonications equivalent to those applied during the ARFI experiment. In this participant (as in others), no detectable temperature rise was present at the target during ultrasound pulses with dose identical to that in the ARFI experiment. The simulated derated intensity values and temperature rise for the other participants are (15.7 W/cm^2^, 0.1°C), (8.7 W/cm^2^, 0.2°C) and (16.4 W/cm^2^, 0.1°C). We confirmed in phantoms that temperature rises of 0.1°C are detectable with our MR-thermometry protocol, ruling out the possibility that the lack of a temperature rise is due to inadequate SNR. Using the same ARFI sonication parameters in a soft tissue phantom resulted in 0.21°C temperature rise at the focal spot. This measurement indicates that ultrasound bioheat simulations may overestimate tissue heating at least in certain situations. Additionally, these data confirm that ARFI signals can be robustly measured in the human brain in situ with no detectable temperature rise at the target.

**Figure 4.**
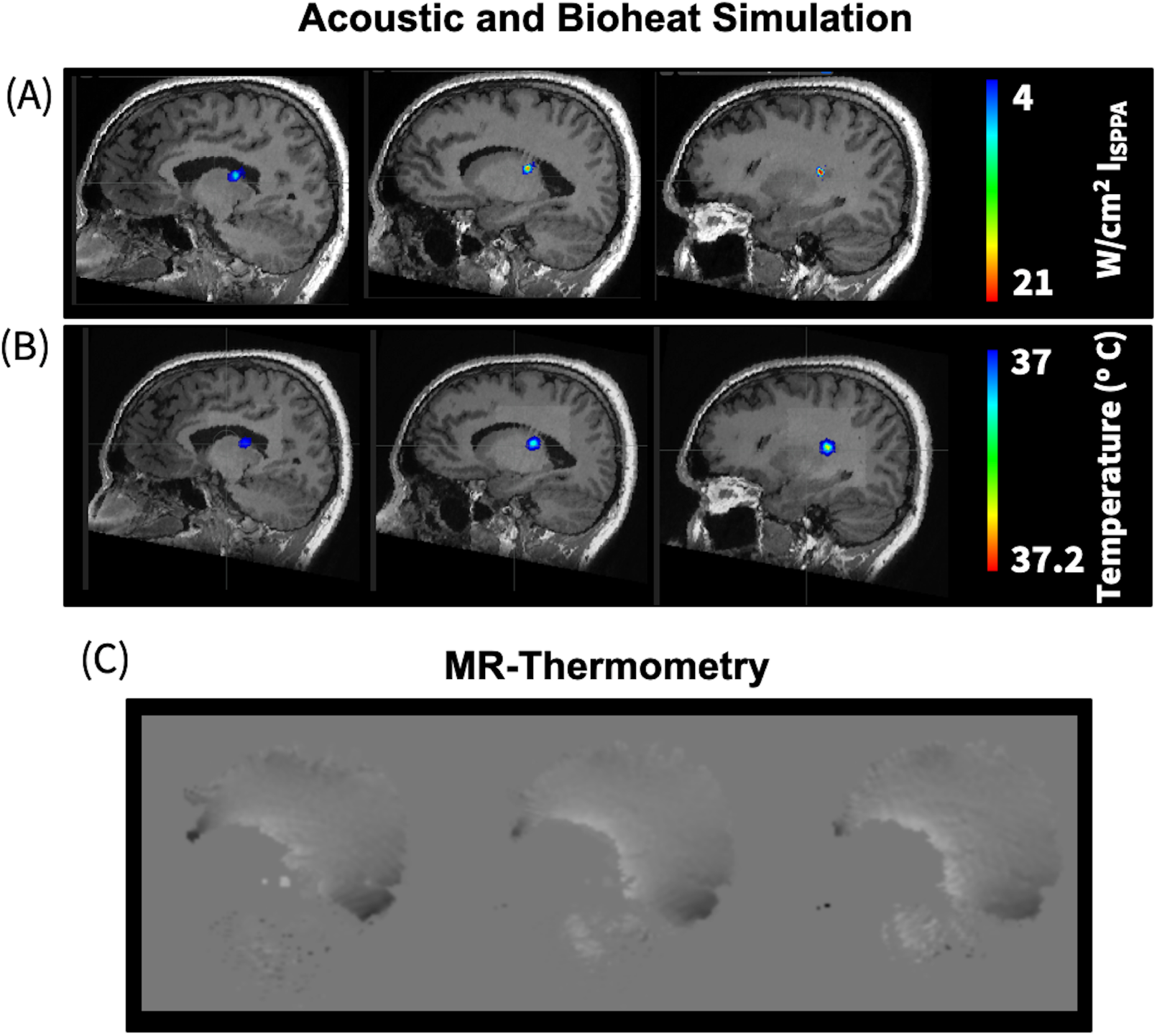
Comparing acoustic simulation and MR-thermometry. This figure shows representative data of the simulated acoustic intensity (A), simulated temperature rise (B) and MR-thermometry (C) for the most transmissive skull amongst participants in this study. The simulated temperature rise for this participant at the focus is < 0.5°C. However, MR-thermometry shows no temperature rise, which is consistent with the cooling effect of the brain’s vasculature (spline interpolation was applied to MR-thermometry images for visualization).

### MR-ARFI in Human

We next attempted to measure ARFI signals in the human brain in situ. As shown in Figure 5, a 500 kHz 4 element annular array (Neurofus System, UK) was coupled to the temporal bone of a human participant. During spiral scans, a free field value of 1.48 MPa (I_SPPA_ = 68 W/cm^2^) at 60 mm was applied. The derated pressure at the focus was estimated as 0.83 MPa (Attali et al, 2022^24^ for skull attenuation and 0.5dB/cm/MHz for brain attenuation). Panel A and B show the planning MRI and the 3D reconstruction of the MR-ARFI in the brain, respectively. Panel C demonstrates t-score heatmap of the ARFI spot overlaid on T1-weighted image in coronal, sagittal and horizontal views. The maximum displacement for this subject was calculated as 1.47 microns.

**Figure 5.**
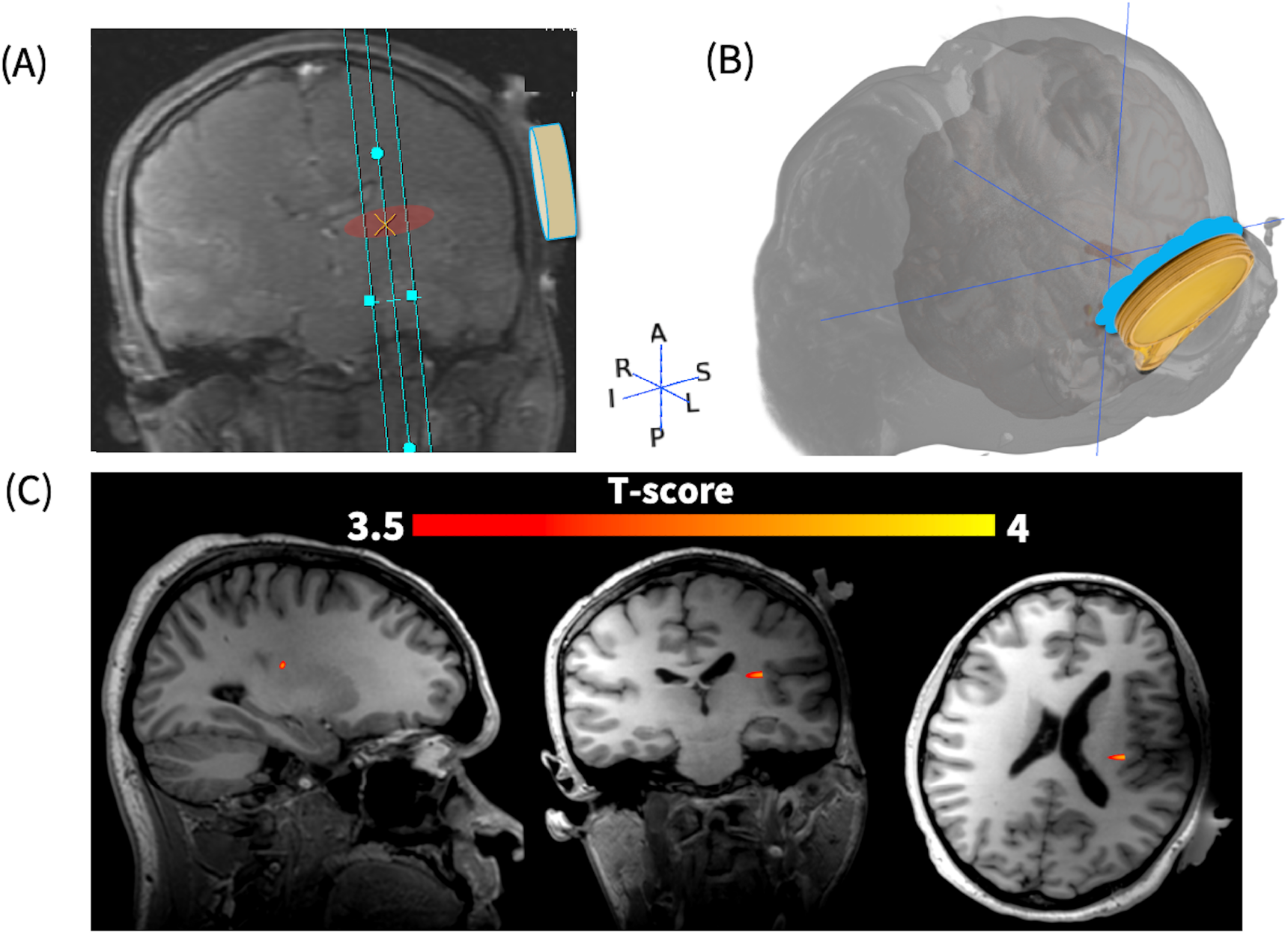
MR-ARFI in human. The planning MRI using three slices centered at 60mm from the transducer (A). 3D rendering of the ARFI spot with the transducer coupled to the left temporal bone (B). Heatmap of the ARFI spot (t-score value) overlaid on T1-weighted image in three planes (C).

We performed ARFI experiments in 5 participants. ARFI maps are shown for 4 participants in Fig. 6A. No ARFI was observed in the fifth participant (not shown), despite experiment repetition with that participant on a different day. Note that there was significant variability in the signal from participant to participant. ARFI T-scores reached significance at p < 0.05 in S1, S3, and S4 with T at least 1.7. In S2, the ARFI signal was observable at the expected focal spot, but the T-score did not reach significance (p>0.28). The variability in measured ARFI signal across participants suggests that different amounts of ultrasound pressure were reaching the target, despite the fact that experimental conditions including transducer positioning, coupling strategy, ultrasound acoustic pressure, sonication protocols, etc. were identical across all conditions. We expect that the differences across participants are primarily due to variations in skull attenuation, which is the primary determinant of ultrasound propagation in brain and is highly variable across participants. Other factors include differences in brain stiffness, hair thickness, brain motion, and variability in the quality of transducer coupling. To assess if differences in skull anatomy could predict differences in ARFI signal across participants, we simulated ultrasound propagation in biophysically realistic simulations using ZTE pseudo-CTs acquired from each participant. Simulated intensity varied significantly from participant to participant (8.7 to 23.5 W/cm^2^). Interestingly the simulated intensity only weakly predicted the ARFI measurement (Fig. 6B, scatter plot of ARFI peak signal, t-score, vs simulated intensity), indicating that differences in simulation accuracy, brain stiffness, brain motion, hair thickness, and/or variability in transducer coupling play a major role in the measured value of ARFI. Given that ARF is the strongest candidate mechanism of ultrasound neuromodulation effects, the fact that simulations do not predict ARFI well emphasizes further how important it is to measure ARFI to be able to determine dosing for each experimental participant.

**Figure 6.**
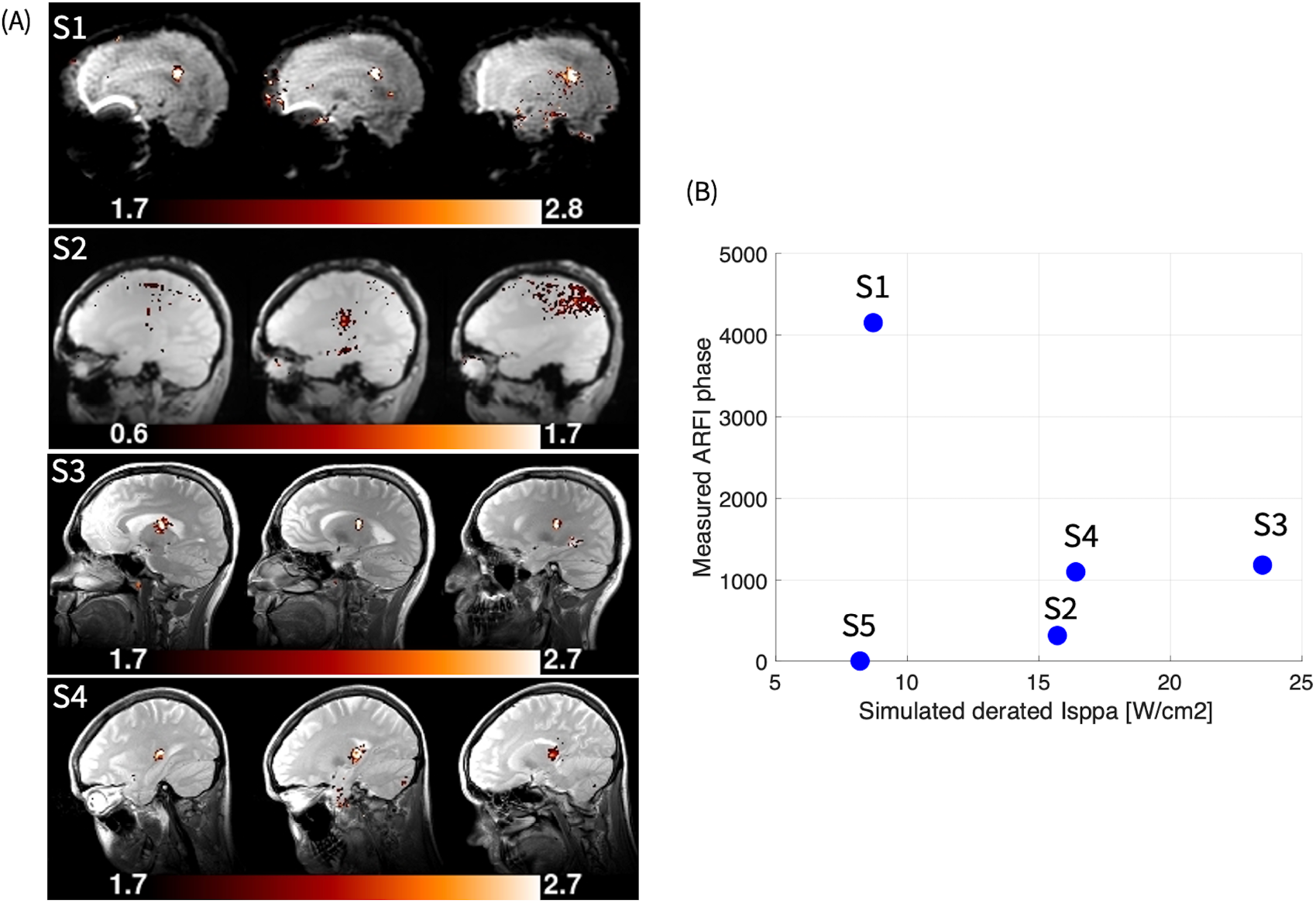
MR-ARFI in human participants. T-score maps of 4 participants; a minimum T-score of 1.7 corresponds to a statistical probability of p < 0.05, uncorrected for multiple comparisons (A). Peak ARFI phase measurements as a function of simulated derated intensity at the focal spot (B).

## Discussion

Here we demonstrate that ARFI can be effectively and efficiently used to 1) confirm the location of the focus, and 2) roughly estimate ultrasound dose at the target, at ultrasound intensities that are less than half of what has previously been reported and that do not produce significant tissue heating (confirmed both by simulation and MR-thermometry). The acoustic parameters used for ARFI in this report are within the safety range of ITRUSST safety consensus^25^. This work provides the transcranial focused ultrasound community with an extremely powerful tool to give ground truth on 1) where in the brain focused ultrasound is delivered (spatial precision), as well as 2) how much ultrasound is delivered (exposure precision).

There is a consensus in the field that reports of highly variable transcranial focused ultrasound effects in published and ongoing studies could be due in large part to uncontrolled variability in the accuracy of targeting^29^. The targeting of transcranial ultrasound depends on simulations which have inherent imprecisions, and both neuronavigated and in-scanner beam positioning solutions require substantial technical skill and are therefore highly operator-dependent. Even a small 5-10^°^ deviation from the desired transducer angle will lead to a complete miss of the target in small deep structures that are the most likely targets of focused ultrasound^30^. ARFI allows experimenters to confirm the focal spot at the desired site, and to reposition the focus (either by manual repositioning of the transducer or electronic beam steering for phase arrays) if the focus is not at the desired location.

Inconsistent ultrasound dosing is also likely a major contributor to variability of reported transcranial ultrasound neuromodulation effects. Although simulations help to an extent, some attenuating properties of the skull (e.g. microporosity) are not captured by ZTE or CT and therefore are not captured by simulation^31,32^. Furthermore, there are practical experimental factors such as quality of coupling, transducer positioning, transducer functioning, and neuronavigation tracking that are not taken into account by simulation yet can have a large effect on the ultrasound dose at the target. The measured ARFI signal accounts for all of these factors and gives a rough ground truth estimate of how much ultrasound is delivered to the target. The data that we report here are T-score statistical parametric maps, which can be converted to an average microscopic displacement in microns (Eq. 2). This knowledge combined with multifrequency MR elastography and tissue stiffness modeling may result in more accurate estimation of *in vivo* acoustic intensity^33^. It is reasonable to predict that ARFI could eventually provide a gold-standard unconfounded measure of individualized in situ ultrasound dose. This will in turn allow a more robust determination of how ultrasound dose relates to neuromodulation effects, which is poorly understood.

The key advance in this paper is our demonstration that the timeseries single-shot MR-ARFI method introduced here has substantial advantages over 2DFT methods, with reduced sensitivity to interference from bulk and pulsatile brain motion. Bulk motion of the head can cause large artifacts in 2DFT ARFI methods because the motion occurs over acquisition of phase-encoding lines of k-space, which takes many seconds or minutes. By contrast, each image of the complex time series single-shot method introduced in this study takes less than 17 ms to acquire, which eliminates intra-image bulk motion effects, while temporal averaging over the time series regains SNR. Furthermore, the short spiral readout (compared with 2x longer EPI readout) reduces susceptibility-drop which increases stability against brain pulsatility^16,34^. Collecting a time series allows all the processing steps of BOLD fMRI to be exploited for low-SNR ARF phase data. Thus, our processing uses a GLM instead of simple subtraction, which allows the incorporation of nuisance regressors to 1) reduce bulk motion with a regressor made from whole-head phase shifts; 2) cycle the ultrasound on and off in multiple blocks to eliminate phase drift while enhancing sensitivity to ARF displacement and reduce temperature rise; and 3) account for physiological pulsation from cardiac and respiratory functions using fMRI-standard processing methods that utilize simultaneously collected cardiac and respiratory data. It has been observed that 2DFT acquisitions with repeated bipolar gradients are more robust against bulk motion and eddy currents compared to inverted bipolar gradients^14,15^. However, using time-series methods with physiological and bulk motion correction, allows inverted bipolar gradients to provide higher ARFI SNR with reduced artifacts.

The time series approach allows for a reduction in ultrasound exposure to 15% that in our previous animal study (Mohammadjavadi et al, 2022)^8^, while improving motion robustness with reduced scan time (80s vs. 205s). The ultrasound parameters used in this study are within the ITRUSST safety recommendations^25^. Simulations estimated at most a 0.2° C temperature rise at the focus, and we did not observe any temperature rise by MR thermometry in our preparation, likely due to the noise limit of in vivo MR-thermometry (∼0.1° C). Using the same sonication parameters in a soft tissue mimicking phantom resulted in maximum 0.21° C temperature rise at the focal spot.

MR-ARFI is promising for guiding many focused ultrasound applications. First, it can noninvasively visualize and verify focal spot without any significant temperature rise. Second, as shown for example in Mohammadjavadi et al 2022, MR-ARFI might be useful to establish a correlation with physiological measures, thus providing a way to establish dose response curve. The method introduced in this study may lead to other novel applications of MR acoustic radiation force imaging.

## CONCLUSION

We demonstrated that the timeseries single-shot MR-ARFI method has substantial advantages over 2DFT methods, by virtue of its reduced sensitivity to interference from bulk and pulsatile brain motion, a reduction in ultrasound exposure (acoustic pressures <0.7 MPa), and reduced scan time (80 seconds per measurement). Future human studies implementing MR-ARFI will be able to achieve more reliable targeting and dosage of low intensity transcranial focused ultrasound.

## Acknowledgments

The authors would like to thank the study participants.

This work was supported by NIH MH131684 and NIH EB032743.

## Data availability statement

The data that support the findings of this study are available from the corresponding author upon reasonable request.

