## Supplemental for "Optimization of MR-ARFI for Human Transcranial Focused Ultrasound"

| | Manufacturer | Centre Frequency | Radius of Curvature | Aperture Diameter | Number Elements | Element Distribution |
|---|---|---|---|---|---|---|
| **Transducer and drive system parameters** | NeuroFUS® (CTX-500) | 500 kHz | 64mm spherical radius | 64 mm | 4 | annular array |

| | Spatial-Peak Pressure Amplitude | Axial Position Spatial-Peak Pressure | Axial - 3dB Width | Lateral - 3dB Width | Axial - 6dB Width | Lateral - 6dB Width |
|---|---|---|---|---|---|---|
| **Free Field Pressure Parameters** | 1.48 MPa | 63 mm | 30-40mm | 6 mm | | 8 mm |

**Table S1. Transducer and drive system description, and free-field pressure parameters.** measurements and simulation. Please refer to the figure S1 demonstrating the pressure beam profile.

| | Duration | Ramp Duration | Ramp shape | Repetition Interval | Notes |
|---|---|---|---|---|---|
| **Pulse** | 6ms | 0 | Rectangular | 21ms | Two pulses of 6ms with 15ms in between, repeated for three MR slices with TR=800ms. |
| **Pulse train** | 27ms | 0 | Rectangular | 266.7ms | |
| **Pulse train repeat** | 80s | 0 | Rectangular | | |

**Table S2_1. Pulse timing parameters for ARFI.** Pulse parameters for MR-ARFI and MR-thermometry experiments. In total there were 50 sonication during the trial, 25 off, 25 ultrasound pulses, 25 off, 25 ultrasound pulses. (see methods, Figure 1). Note that in subject 4 we used two 7ms pulses of ultrasound for ARFI instead of 6ms.

| | Duration | Ramp Duration | Ramp shape | Repetition Interval | Notes |
|---|---|---|---|---|---|
| **Pulse** | 12ms | 0 | Rectangular | 12ms | |
| **Pulse train** | 12ms | 0 | Rectangular | 266.7ms | |
| **Pulse train repeat** | 80s | 0 | Rectangular | | |

**Table S2_2. Pulse timing parameters for thermometry.** Pulse parameters for MR-thermometry experiments. Note that in subject 4 we used 14ms instead of 12ms.

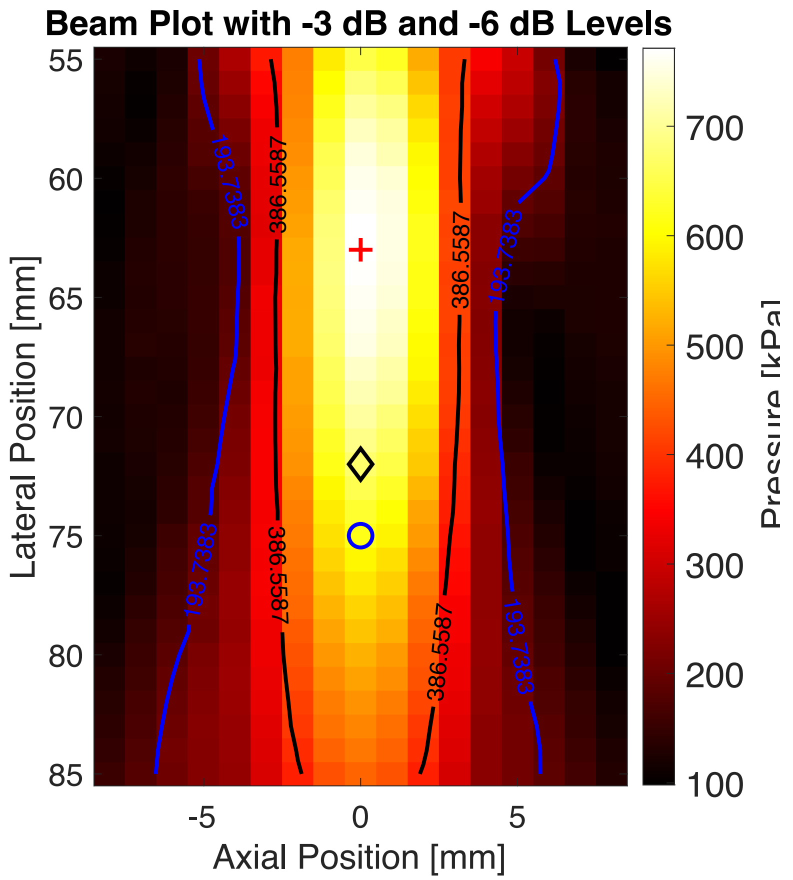


**Figure S1. Pressure field at the focal spot.** Focusing on-axis at 65 mm away from the transducer exit plane. Free field pressure parameters are shown as: black contour is -3 dB focal region relative to maximum pressure, blue contour is -6dB region relative to maximum pressure and the red cross is the axial position of the location of spatial-peak pressure. Note that the maximum free field pressure during the experiment was 1.48 MPa, however the measurement was performed at 800kPa pressure values ($I_{sppa}$ = 20 $W/cm^2$) to avoid damage to the hydrophone's membrane.
